# PARP14 is a writer, reader and eraser of mono-ADP-ribosylation

**DOI:** 10.1101/2023.06.24.546374

**Authors:** Archimede Torretta, Constantinos Chatzicharalampous, Carmen Ebenwaldner, Herwig Schüler

**Affiliations:** Center for Molecular Protein Science (CMPS), Department of Chemistry, Lund University, 22100 Lund, Sweden

**Keywords:** ADP-ribosyltransferase, ADP-ribosylation, hydrolase, macrodomain, poly(ADP-ribose) polymerase (PARP), post-translational modification (PTM), protein domain, signaling

## Abstract

PARP14/BAL2 is a large multidomain enzyme involved in signaling pathways with relevance to cancer, inflammation, and infection. Inhibition of its mono-ADP-ribosylating PARP homology domain and its three ADP-ribosyl binding macro domains has been regarded as a potential means of therapeutic intervention. Macrodomains-2 and -3 are known to stably bind to ADP-ribosylated target proteins; but the function of macrodomain-1 has remained somewhat elusive. Here, we used biochemical assays of ADP-ribosylation levels to characterize PARP14 macrodomain-1 and the homologous macrodomain-1 of PARP9. Our results show that both macrodomains display an ADP-ribosyl glycohydrolase activity that is not directed toward specific protein side chains. PARP14 macrodomain-1 is unable to degrade poly(ADP-ribose), the enzymatic product of PARP1. The F926A mutation of PARP14 and the F244A mutation of PARP9 strongly reduced ADP-ribosyl glycohydrolase activity of the respective macrodomains, suggesting mechanistic homology to the Mac1 domain of the SARS-CoV-2 Nsp3 protein. This study adds two new enzymes to the previously known six human ADP-ribosyl glycohydrolases. Our results have key implications for how PARP14 and PARP9 will be studied and how their functions will be understood.

## Introduction

Intracellular ADP-ribosylation in human physiology is catalyzed by enzymes of the poly(ADP-ribose) polymerase (PARP) family. The name-giving enzyme, PARP1, and its nearest homologues use NAD+ to build chains of ADP-ribose onto amino acid side chains in their target proteins (1). Like the ADP-ribosylating bacterial toxins, however, most PARP family members catalyze mono-ADP-ribosylation (MARylation)(2-4). PARP enzymes appear to MARylate proteins predominantly at serine and carboxylic acid residues, but other acceptor side chains have been identified (5,6). PARP enzymes likely also contribute to MARylation of non-protein substrates (7).

Posttranslational modifications often affect cellular processes in part by acting as anchoring points for binder domains and MARylation is no exception. The main recognition module for MARylation is the macrodomain, a ~190 residue module found in all kingdoms of life that can bind and in many cases hydrolyze various ADP-ribosyl adducts and/or metabolites (8-11). The human genome encodes 11 macrodomain containing proteins (11). The macrodomains of histone variants MacroH2A1 and -2, the chromatin remodeler CHD1L/ALC1, the Ganglioside-induced differentiation-associated protein 2 (GDAP2) and the MARylating PARP family members PARP9, -14 and -15 are thought to be ADP-ribosyl binding domains. By contrast, MacroD1 and MacroD2 (12,13), TARG1 (14,15) and PARG (16) macro-domains have ADP-ribosyl hydrolase activity; and TARG1 can process both proteins and nucleic acids as substrates (17). The three PARP enzymes stand out in that they contain two (PARP9, PARP15) and three (PARP14) macro-domains in series (11).

PARP14 was first described as B-cell aggressive lymphoma protein-2 (BAL2), along with PARP9/BAL1 and PARP15/ BAL3, as frequently overexpressed in fatal high-risk lymphoma (18,19). PARP14 is involved in multiple signaling pathways linked to cancer, inflammation, and infection (20-23) and pharmacological modulation of PARP14 activities may prove as a suitable therapeutic strategy. In all mammals, PARP14 is a >1800 residue multidomain protein. The N-terminal half of PARP14 consists of a series of nucleic acid binding RRM- and KH domains. The seventh of the eight KH domains is split in two by insertion of the three macrodomains. The C-terminus consists of a WWE domain followed by the MARylating PARP homology domain (or mART domain). In vitro studies show that the isolated PARP homology domain has efficient auto-MA-Rylation activity (24), modifies PARP14 macrodomains-2 and -3 efficiently (25) and is unable to modify RNA (26). In cells, endogenous PARP14 has been found to be modified on two histidine and three tyrosine residues, but it is not known whether these modifications are a product of auto- or trans-modification. (27) Overexpression of PARP14 variants in cells combined with a chemical genetics approach and subsequent target identification suggested that PARP14 can modify over 110 proteins including PARP1 and PARP13, and a gene ontology analysis corresponded well with the previously suggested RNA regulation related functions of PARP14 (28). The same study showed that PARP14 primarily modified carboxylic acid side chains (28) and this has also been suggested based on MS analysis of peptides ADP-ribosylated in vitro by the PARP14 catalytic domain (29).

It is undebated that PARP14 macrodomains-2 and -3 behave as expected of ADP-ribosyl binder domains in vitro. They form stable interactions with auto-MARylated protein (30) and this has been employed to construct a recognition module, a tandem macrodomain-Fc fusion for use as a research tool (31). They also lack the active-site side chains that have been implicated in catalysis in hydrolytically active macrodomains (11). By contrast, the ADP-ribosyl binding of macrodomain-1 has been more difficult to establish. Macrodomain-1 has a documented low affinity for free ADP-ribose (32). Moreover, neither PARP14 macro-domain-1 nor the homologous PARP9 macrodomain-1 interacted stably with any of several auto-MARylated PARP catalytic domains examined (30). This finding might be explained by an ADP-ribosyl glycohydrolase activity of macrodomain-1, and there is indeed published evidence for such activity (25,32). Two studies (13,33) have highlighted the conservation of residues potentially involved in ADP-ribosyl glycohydrolase activity between PARP14 macrodomain-1 and the first macrodomain of the SARS-CoV Nsp3 protein, a macrodomain that had already been shown to be catalytically active (34,35). Following the advent of SARS-CoV-2, the first macrodomain of the SARS-CoV-2 Nsp3 protein (often called Mac1) was also shown to possess ADP-ribosyl glycohydrolase activity (25,36) and its homology with PARP14 macrodomain-1 was pointed out as well (25,37). Despite all this evidence, the question of whether the N-terminal macrodomains of PARP14 and PARP9 have ADP-ribosyl glycohydrolase activity has not been explicitly addressed by experiment.

Here we show that PARP14 macrodomain-1, but neither macrodomain-2 nor macrodomain-3, can remove MAR generated by the mART domains of PARP14 and PARP10 in vitro. This activity is more efficient when macrodomain-1 is presented within a protein construct containing all three macrodomains. PARP14 macrodomain-1 lacks amino acid selectivity and is unable to degrade poly(ADP-ribose). As expected based on sequence homology, PARP9 mac-rodomain-1 also possesses glycohydrolase activity. Several compounds recently found to inhibit the SARS-CoV-2 Nsp3 Mac1 showed no effect on either PARP14 or PARP9 macrodomain-1 activity. The F926A mutant of PARP14 and the F244A mutant of PARP9, while being able to bind free ADP-ribose, have strongly reduced ADP-ribosyl glycohydrolase activities.

## Experimental Procedures

### Bioinformatics

Human PARP14 (Uniprot: Q460N5) and PARP9 (Q8IXQ6) macrodomains, macroD2 (A1Z1Q3) and macrodomain-1 of SARS-CoV-2 Nsp3 (P0DTC1) sequences were aligned by Clustal Omega and secondary structural elements were added to the alignment using ESPript (38). The phylogenetic tree was calculated starting from the Clustal Omega alignment using the neighbour-joining algorithm of MEGA11 (39). Structural superimpositions and RMSD calculations between Cα atom pairs were performed with PyMOL (Schrödinger, Inc.). Molecular graphics images were prepared with ChimeraX (40).

### Materials

All chemicals and reagents were purchased from Merck Life Science unless otherwise specified, and all chromatography media were purchased from Cytiva.

### Molecular cloning

Human PARP14^L1449-K1801^ construct was obtained by sub-cloning the genetic sequence, optimized for expression in E. coli, in pNIC28. PARP14 macrodomain-1^S794-D984^, macrodomain-2^A994-S1196^, macrodomain-1+2^G784-S1196^, macro-domain-3^F1208-G1388^, macrodomain-2+3^A994-G1388^, and macro-domain-1+2+3^G784-S1393^ constructs, (32) human PARP9 mac-rodomain-1^G67-K263^ and PARP10^N819-V1007^ constructs (30), ARH1 (41) and ARH3, uman PARP1^K662-T1011^, HPF1, macroD2, and Af1521-GFP fusion constructs (42) were previously described. The human PARP14 macrodomain-1 F926A mutant and the human PARP9 macrodomain-1 F244A mutant were obtained by subcloning the E. coli codon optimized synthetic cDNA into pET151-D-TOPO using the respective sequences of the wild-type macrodomain-1 as templates (GeneArt, Thermo Fisher Scientific). Synthetic cDNA encoding Clostridium botulinum C2I toxin subunit was sub-cloned in pET28 as described (43).

### Protein Purification

All protein constructs were transformed and expressed in E. coli BL21(DE3)T1R (Merck Life Science). Cells were co-transformed with the pRARE2 plasmid (Protein Science Facility, Karolinska Institutet, Stockholm, Sweden) unless codon-optimized cDNAs were used. Overnight pre-cultures in Terrific Broth (TB) media supplemented with relevant antibiotics (34 µg/mL chloramphenicol; 50 µg/mL kanamycin; 100 µg/mL ampicillin) were inoculated in fresh TB media supplemented with antibiotics and cells were grown at 37ºC using a LEX bioreactor (Epiphyte3). When OD_600_ reached 2.0, recombinant protein expression was induced with 0.5 mM IPTG at 18ºC for 16 hours. Cells were harvested by centrifugation at 4500xg and resuspended in 50 mM HEPES pH 7.5, 500 mM NaCl, 10 %v/v glycerol, 10 mM imidazole, 0.5 mM TCEP buffer supplemented with EDTA-free protease inhibitor cocktail and benzonase. The resuspended pellets were lysed by sonication and the cell lysate was clarified by centrifugation at 24000xg. The lysate was filtered (0.45 µm) and loaded onto HiTrap chelating HP columns charged with Co^2+^ ions for Immobilized Metal Affinity Chromatography (IMAC). The loaded columns were washed with 50 mM HEPES pH 7.5, 500 mM NaCl, 10 %v/v glycerol, 10 mM imidazole, 0.5 mM TCEP buffer (10 column volumes) and elution was performed with a step gradient of 260 mM imidazole in 50 mM HEPES pH 7.5, 300 mM NaCl, 10 %v/v glycerol, 0.5 mM TCEP buffer. IMAC fractions were analyzed by SDS-PAGE and fractions containing target protein were pooled and further purified by size-exclusion chromatography (SEC) using a Super-dex-75 column (16/60) equilibrated with 50 mM HEPES pH 7.5, 300 mM NaCl, 10 %v/v glycerol, 0.5 mM TCEP buffer. The purified proteins were concentrated by ultrafiltration and stored at -80ºC. Bovine cytosolic γ-actin was purified from calf thymus as described (44).

### Western blot analysis

PARP14^L1449-K1801^ and PARP10^N819-V1007^ automodification reactions were performed in the presence of 100 µM NAD+ (10% biotinylated; BPS Bioscience) and with 1 µM of enzyme in reaction buffer (50 mM HEPES pH 7.5, 100 mM NaCl, 0.2 mM TCEP, 4 mM MgCl2, 0.1 mM EDTA). Master reactions were incubated at room temperature for 1 hour before the reactions were stopped by addition of 100 µM of RBN012759 (MedChemExpress) for PARP14, or PJ34 (Merck Life Science) for PARP10. Negative controls contained PARP inhibitors from the onset. An aliquot from the starting master reaction was supplemented with SDS-containing loading buffer and inactivated for 4 minutes at 95ºC, while the remaining volume was incubated with the different PARP14 macrodomain constructs or PARP9 macrodomain-1 at a final concentration of either 3 or 10 µM. Aliquots were taken at intervals or after 4 hours and inactivated by heating with SDS-PAGE loading buffer. The collected aliquots (technical duplicates) and the SeeBlue Plus2 PreStained Protein Standard (Thermo Fisher Scientific) were resolved by SDS-PAGE and transferred to a Immobilon-FL PVDF membrane by wet transfer. The membrane was incubated for 30 minutes each with 1% BSA in TBST buffer followed by Streptavidin-HRP (Thermo Fisher Scientific) in TBST buffer. Development was performed using the Clarity™ Western ECL Substrate (Bio-Rad) and images were taken with the ChemiDoc Imaging System (Bio-Rad). Chemiluminescence was analyzed with the Image Lab software (Bio-Rad) and normalized intensities were obtained by dividing the band intensity measured at a given time by the one at time zero and averaging over the duplicates. Results were plotted with GraphPad Prism 9.

### ADP-ribosylation reactions for the plate-based MacroGreen overlay assay

For PARP14 and PARP10 automodification reactions, 1 µM of enzyme was incubated with 100 µM of NAD+ in 50 mM HEPES pH 7.5, 100 mM NaCl, 0.2 mM TCEP, 4 mM MgCl2, 0.1 mM EDTA reaction buffer. Reactions were incubated for 30 minutes at 37ºC and then stopped by adding RBN012759 or PJ34 at a final concentration of 100 µM. For auto-PARylation of PARP1, 1 µM of enzyme was incubated with 1 mM of NAD+ in 50 mM HEPES pH 7.5, 100 mM NaCl, 0.2 mM TCEP, 4 mM MgCl2 reaction buffer, while, to obtain serine ADP-ribosylation, 1 µM of PARP1 was incubated with 1 µM of HPF1 with 1 mM of NAD+ in the same reaction buffer. Reactions were incubated for 30 minutes at 37ºC and then stopped by adding Talazoparib (Selleck Chemicals) at a final concentration of 1 µM. To produce arginine ADP-ribosylation, 20 nM of Clostridium botulinum C2I toxin was incubated with 10 µM of γ-actin in 50 mM HEPES pH 7.5, 100 mM NaCl, 0.2 mM TCEP, 4 mM MgCl2 reaction buffer. Reactions were incubated for 30 minutes at 37ºC.

### Quantitative assay of ADP-ribosylation levels using MacroGreen overlay

50 µL of MAR- or PARylated samples were dispensed in each well of a MaxiSorpTM plate (Nunc) and incubated at room temperature for 30 minutes with shaking at 250 rpm. Unbound proteins were removed by discarding the liquid in the wells and the plate was washed with 150 µL per well of 50 mM HEPES pH 7.5, 100 mM NaCl, 0.2 mM TCEP, 4 mM MgCl2, 0.1 mM EDTA buffer for three times. 150 µL per well of 1% BSA in TBST were added and the plates were incubated for 5 minutes with shaking at 250 rpm. The wells were washed with 150 µL of 50 mM HEPES pH 7.5, 100 mM NaCl, 0.2 mM TCEP, 4 mM MgCl2, 0.1 mM EDTA buffer for three times. 50 µL per well of solutions of different macrodomains at a final concentration of 6 µM were added to the wells, except for positive control (omission of macrodomain), negative control (omission of NAD+) and blank, where 50 µL of 50 mM HEPES pH 7.5, 100 mM NaCl, 0.2 mM TCEP, 4 mM MgCl2, 0.1 mM EDTA buffer was added. The plates were incubated at 37ºC for 1 hour in a thermomixer with shaking at 250 rpm. Wells where rinsed and washed once with 150 µL per well of 50 mM HEPES pH 7.5, 100 mM NaCl, 0.2 mM TCEP, 4 mM MgCl2, 0.1 mM EDTA buffer, then wells containing macrodomains were incubated with 100 µL of 1 mM ADP-ribose at room temperature for 10 minutes with shaking at 250 rpm, to remove bound macrodomains prior to detection of ADP-ribosylation. Plates were rinsed again, washed with 150 µL per well of TBST buffer for three times, blocked with 150 µL per well of 1% BSA in TBST buffer for 5 minutes at room temperature with shaking at 250 rpm and once with 150 µL per well of TBST buffer. 50 µL per well of MacroGreen, a multi-site mutant of Af1521 fused to green fluorescent protein, (42) were added and the plates were incubated for 5 minutes at room temperature in the thermomixer with shaking at 250 rpm. Finally, 150 µL per well of TBST where used to wash the plates and then GFP-fluorescence measurements were performed in the presence of 150 µL per well of TBST with the CLARIOstar plate reader (BMG Labtech) using a 470-15 nm excitation filter and a 515-20 nm emission filter. Measurements were performed in quadruplicate. The average fluorescence intensities from the negative control and the blank were subtracted from each well and the results were normalized with the average intensity from the positive control. Graphs were plotted and statistical analyses (One-way ANOVA test) were done using GraphPad Prism 9 (n=4; significance levels: * 0.0332; ** 0.0021; *** 0.0002; **** 0.0001; all significance levels refer to the control values in the first columns).

### Thermal stability and ligand induced Tm shift analysis by differential scanning fluorimetry

Melting point (T_m_) assays were performed in white 96-well PCR plates (Bio-Rad #MLL9651). Every well contained a 25 µL solution of 0.4 mg/mL of enzyme (with or without 2 mM of ADP-ribose) and SyproOrange (ThermoFisher; at 1:5000 dilution) in 50 mM HEPES pH 7.5, 300 mM NaCl, 10 %v/v glycerol, 0.5 mM TCEP buffer. Plates were incubated at 20 ºC for 10 seconds and then the temperature was increased by 1 ºC/min up to 95 ºC. Fluorescence signals were measured with the CFX96 Touch Real-Time PCR Detection System (Bio-Rad) and data analysis was performed with the CFX Manager software (version 3.1, Bio-Rad).

## Results and Discussion

### PARP14 macrodomain-1 has ADP-ribosyl glycohydrolase activity in vitro

Several lines of evidence, summarized above, suggested that PARP14 macrodomain-1 may possess ADP-ribosyl glycohydrolase activity. We tested whether this was indeed the case by allowing auto-MARylation of the catalytic domain construct PARP14^L1449-K1801^ in the presence of biotin-NAD+, followed by incubation with various PARP14 macrodomain combinations and detection of biotinylated substrate levels by Western blotting. Fig. 1A shows that macrodomain-1 efficiently removes automodification from the PARP14 catalytic domain. By contrast, neither macro-domain-2 nor macrodomain-3 affected the levels of automodified catalytic domain (Supplementary Materials, Fig. S1B,C). When the macrodomains were presented within constructs spanning several domains, an apparent increase in catalytic efficiency was notable for macrodomain-1, while macrodomains-2 and -3 remained inactive (Fig. 1B-D and Supplementary Materials, Fig. S1). The same results were obtained using the catalytic domain construct PARP10^N819-V1007^ (Figure 1E). This shows that PARP14 macrodomain-1 is an ADP-ribosyl glycohydrolase that is more efficient within the context of macrodomains-2 and -3 (Fig. 1F) representing the physiological arrangement (Fig. 2A).

**Figure 1.**
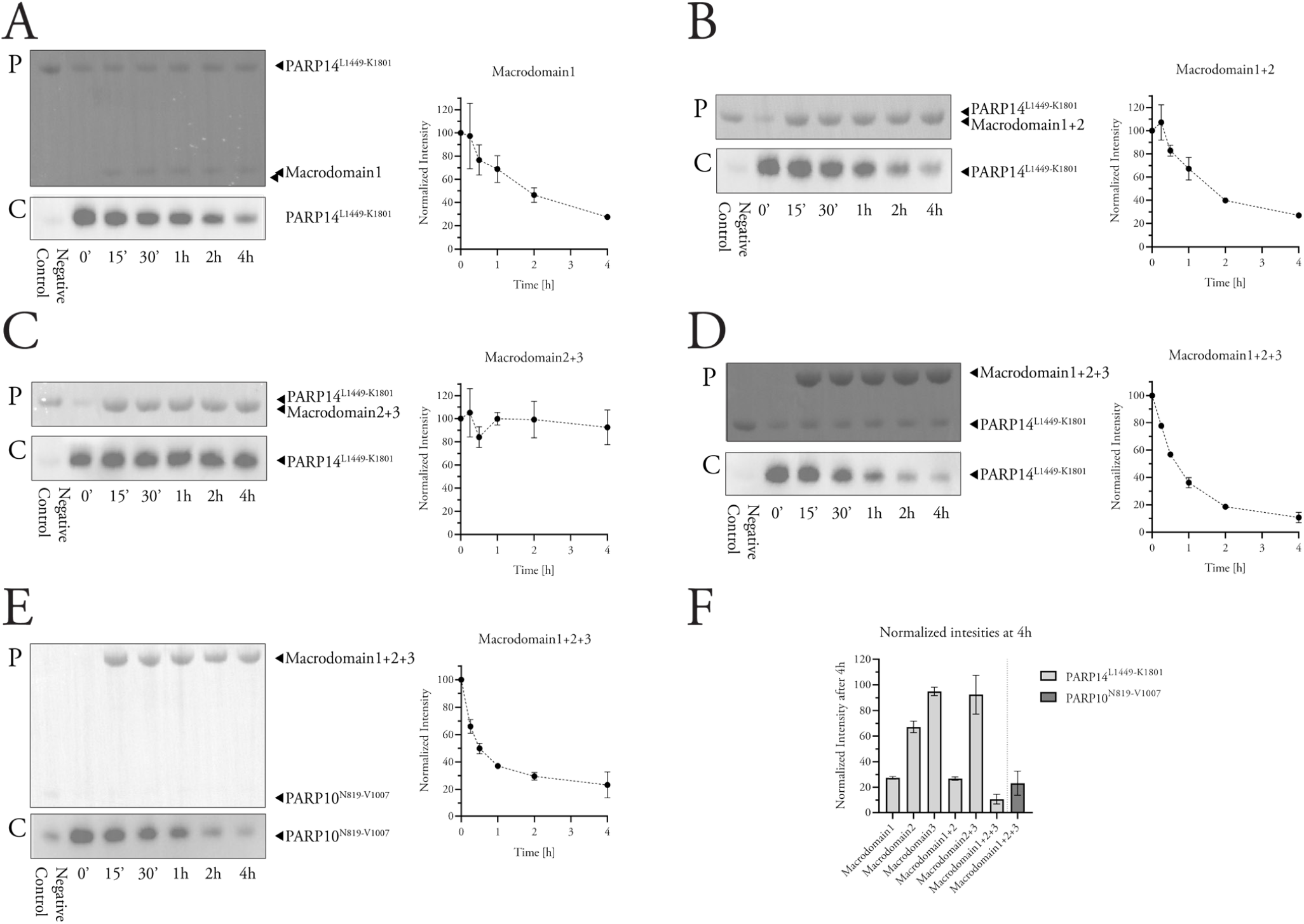
ADP-ribosyl glycohydrolase activity of PARP14 macrodomain-1 measured by Western blotting. (**A-D**) Automodified PARP14 catalytic domain construct was incubated with PARP14 macrodomain-1 (**A**), macrodomain-1+2 (**B**), macrodomain-2+3 (**C**), or macrodomain-1+2+3 (**D**) and the resulting levels of automodification (biotin-ADP-ribosyl) were detected using streptavidin-HRP. (**E**) Automodified PARP10 catalytic domain construct incubated with PARP14 macrodomain 1+2+3 and analyzed as above. Upper panels labeled with “P” show Ponceau S-stained membranes after transfer, while lower panels labeled with “C” show chemiluminescence images. The normalized intensities measured from each experiment are shown to the right (means±S.D. of two experiments). (**H**) Comparison of the normalized intensities (means±S.D. of two experiments) of the last time points of each experiment (**A-E**). Full membranes for these experiments are shown in the Supplementary Materials, Fig. S1).

**Figure 2.**
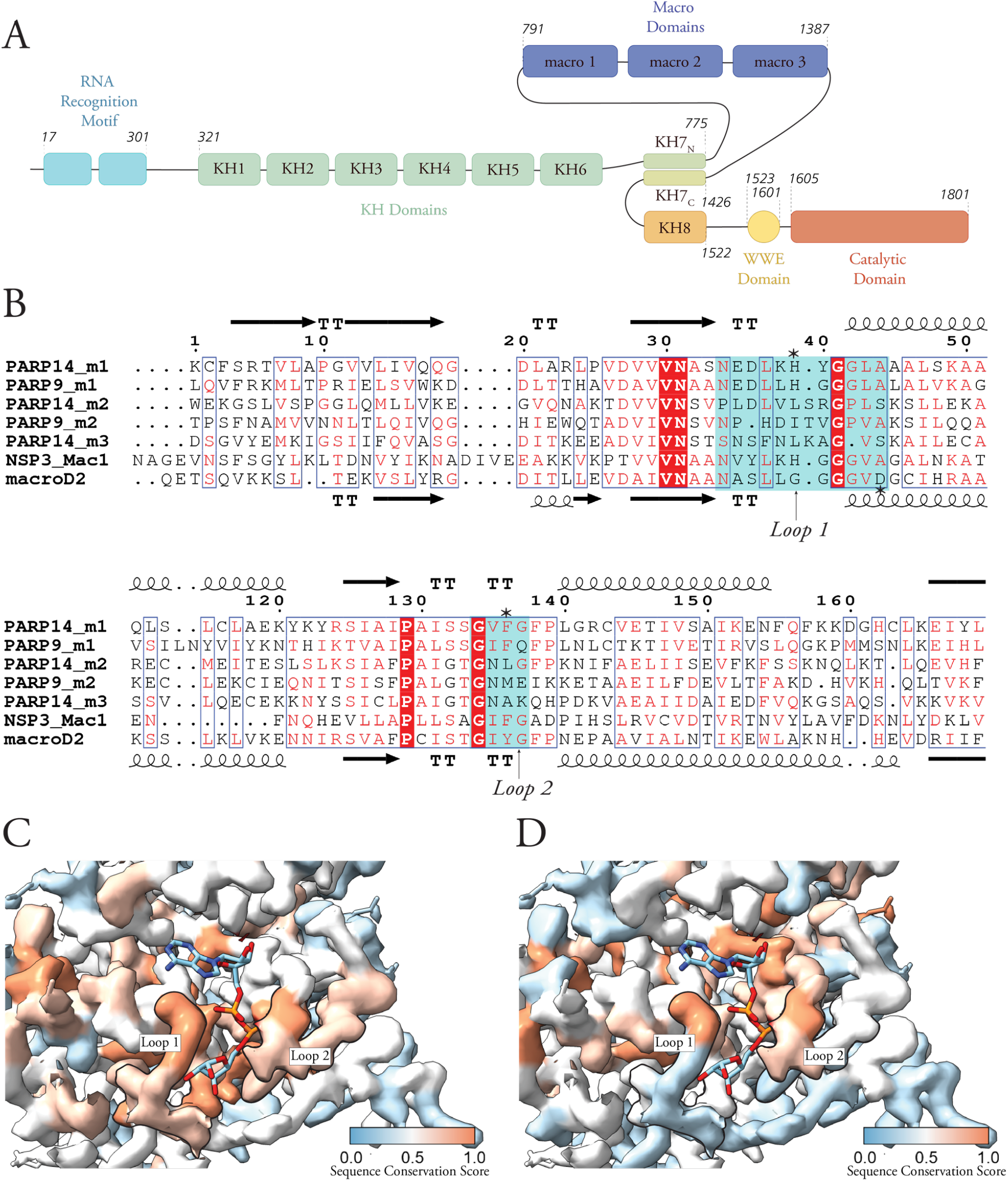
PARP14 domain arrangement and macrodomain evolutionary relationships. (**A**) Schematic illustration of PARP14 domain structure with the multiple RRM and KH domains, three macrodomains, WWE domain, and the PARP-homology ADP-ribosyl transferase domain. (**B**) Sequence alignment of the indicated PARP14 and PARP9 macrodomains, macroD2, and the SARS-COV-2 Nsp3 Mac1. The secondary structures of PARP14 macrodomain-1 (PDB: 3Q6Z) and macroD2 (4IQY) are shown above and below the alignment, respectively. Cyan boxes indicate the positions of loops 1 and 2 and asterisks mark residues referred to in the text. (**C**) Electron density from the crystal structure of PARP14 macrodomain-1 (3Q6Z) colored according to the conservation score calculated by aligning PARP14 macrodomain-1 with the catalytically active PARP9 macrodomain-1 (Alphafold prediction), macroD2 (4IQY), and SARS-COV-2 Mac1 (6WOJ). (**D**) Electron density from the crystal structure of PARP14 macrodomain-1 colored according to the conservation score calculated by aligning PARP14 macrodomain-1 with the ADP-ribosyl binders PARP14 macrodomain-2 (3Q71), PARP14 macrodomain-3 (4ABK), and PARP9 macrodomain-2 (5AIL). Thick black outlines contour the positions of loops 1 and 2. The conservation score is a number between 0 for dissimilar residues, and 1 for identical residues.

### Sequence and structure features of PARP14 macrodomain-1

To identify the sequence features responsible for the ADP-ribosyl glycohydrolase activity of macrodomain-1, we aligned the sequences of the human PARP14 and PARP9 macrodomains, macroD2 and SARS-CoV-2 Nsp3 Mac1 (Fig. 2B). These macrodomains share an average residue identity of approximately 30% and a similarity of 40% (Table S1). All these domains are characterized by an identical globular folding topology, the α/β/α sandwich fold (Fig. S2A). The calculated pair-wise root mean square deviations (RMDS) between Cα positions are generally less than 2 Å, although PARP14 macrodomains-2 and -3 display a somewhat greater difference to the Nsp3 Mac1 (Supplementary Materials, Table S1). Despite the overall structural similarity, the phylogenetic tree resulting from the alignment identified two clades with PARP14 and PARP9 macrodomain-1 closer to the two glycohydrolytically active macrodomains than to the other PARP macrodomains (Fig. S2B), as previously found (33). The macroD-protein related mono(ADP-ribosyl) hydrolases are characterized by conserved sequence motifs, N(6x)GG[V/L/I] and G[V/ I/A][Y/F]G in the two substrate binding loops (loops 1 and 2, also called the catalytic and the diphosphate binding loop) (11). The sequence alignment showed that these two motifs are conserved in PARP14 and PARP9 macrodomains-1, while they are absent in the catalytically inactive macro-domains-2 and -3 (Fig. 2B, boxed in cyan).

Although several residues may make alternative or target specific contributions to catalysis (41), studies on macroD1 and -2 and the SARS-CoV-2 Nsp3 Mac1 indicate that the E or H side chains in the catalytic loop (marked with an asterisk in Fig. 2B) are responsible for the deprotonation of a water molecule that then performs the hydrolysis at the MARylated side chain, while the substrate is held in position by the diphosphate loop (15,45-47). In particular, the role of an aromatic residue (Phe or Tyr; marked by an asterisk in Fig. 2B) in the diphosphate loop has been proposed to be crucial for securing the correct orientation of the ADP-ribose moiety inside the catalytic pocket through steric hindrance and dipole interactions (48). This site is conserved among macroD-like mono(ADP-ribosyl) hydrolases (48). Taken together, our comparison indicates that PARP14 and PARP9 macrodomains-1 possess the sequence features of MAR hydrolases, which clearly distinguishes them from the remaining PARP macrodomains.

Large-scale in silico and in vitro compound screening efforts have recently identified several compounds that inhibit SARS-CoV-2 Nsp3 Mac1 activity (49-51). The sequence and structural similarities between PARP14 and PARP9 macrodomains-1 and SARS-CoV-2 Nsp3 Mac1 prompted us to test three compounds that had been identified as inhibitors of Mac1 (51) for their effect on PARP14 and PARP9 macrodomain-1 activity. While the compounds reduced the melting temperature of PARP14 macrodomain-1, they had no measurable effect on either PARP14 or PARP9 macrodomain-1 activity (Figs. S3 and S4). We conclude that these compounds may form interactions with a thermally unstable (possibly open-pocket) conformation of PARP14, a binding mode that is mechanistically different from their binding to Nsp3 Mac1 (51). In the absence of an effect on enzymatic activity, these compounds are not suitable as starting points for development of PARP14 or PARP9 macrodomain-1 inhibitors.

### Macrodomain-1 target preferences

Having established macrodomain-1 ADP-ribosyl glycohy-drolase activity and its sequence similarity to macroD proteins, we asked whether macrodomain-1 also shared the preference for MARylated carboxylic acid residues (Glu, Asp) attributed to other macrodomain glycohydrolases (41). To this end, we compared macrodomain-1 activity to the activities of macroD2 and of the ADP-ribosylhydrolases ARH1 and ARH3, which are known to be specific for Arg- and Ser-linked modification, respectively (41,52). First, we incubated the panel of our macrodomains and the ADP-ribosylhydrolases with automodified PARP14, as above. Then, we quantified remaining ADP-ribosylation using MacxroGree,n, a GFP-tagged ADP-ribosyl binder, in overlay assays. As shown in Fig. 3A, this experiment confirmed the inactivity of macrodomains-2 and -3 and the efficient removal of PARP14 auto-MARylation by macro-domain-1. This was important to establish to exclude the possibility that the biotinyl on ADP-ribose (used in the experiments shown in Fig. 1) had detrimental effects on macrodomain activities. ARH1 did not reduce auto-MARylation, indicating that PARP14 does not auto-MARylate on Arg side chains. The Ser-specific ARH3 (53) and the Glu/ Asp-specific macroD2 (12), however, both reduced MARy-lation to very low levels (Fig. 3A). These observations might be explained by a scenario where (i) PARP14 auto-MARy-lates on Ser and carboxylic acid side chains to a similar extent; and (ii) ARH3 and macroD2 substrate specificities are not as strict as thought and both can cleave ADP-ribosyl off Ser and carboxylic acid side chains to a similar extent. Alternatively, automodification of an altogether different side chain might be at play. Since PARP14 has been found to be MARylated on His and Tyr side chains (27), our observations suggests that PARP14 auto-MARylates on these side chains and macrodomain-1 as well as both ARH3 and macroD2 are capable of ADP-ribosyl removal from them. Given the different chemistries involved, we believe that our findings are in favour of PARP14 Tyr auto-MARylation and subsequent ADP-ribosyl cleavage off Tyr by all three glycohydrolases.

**Figure 3.**
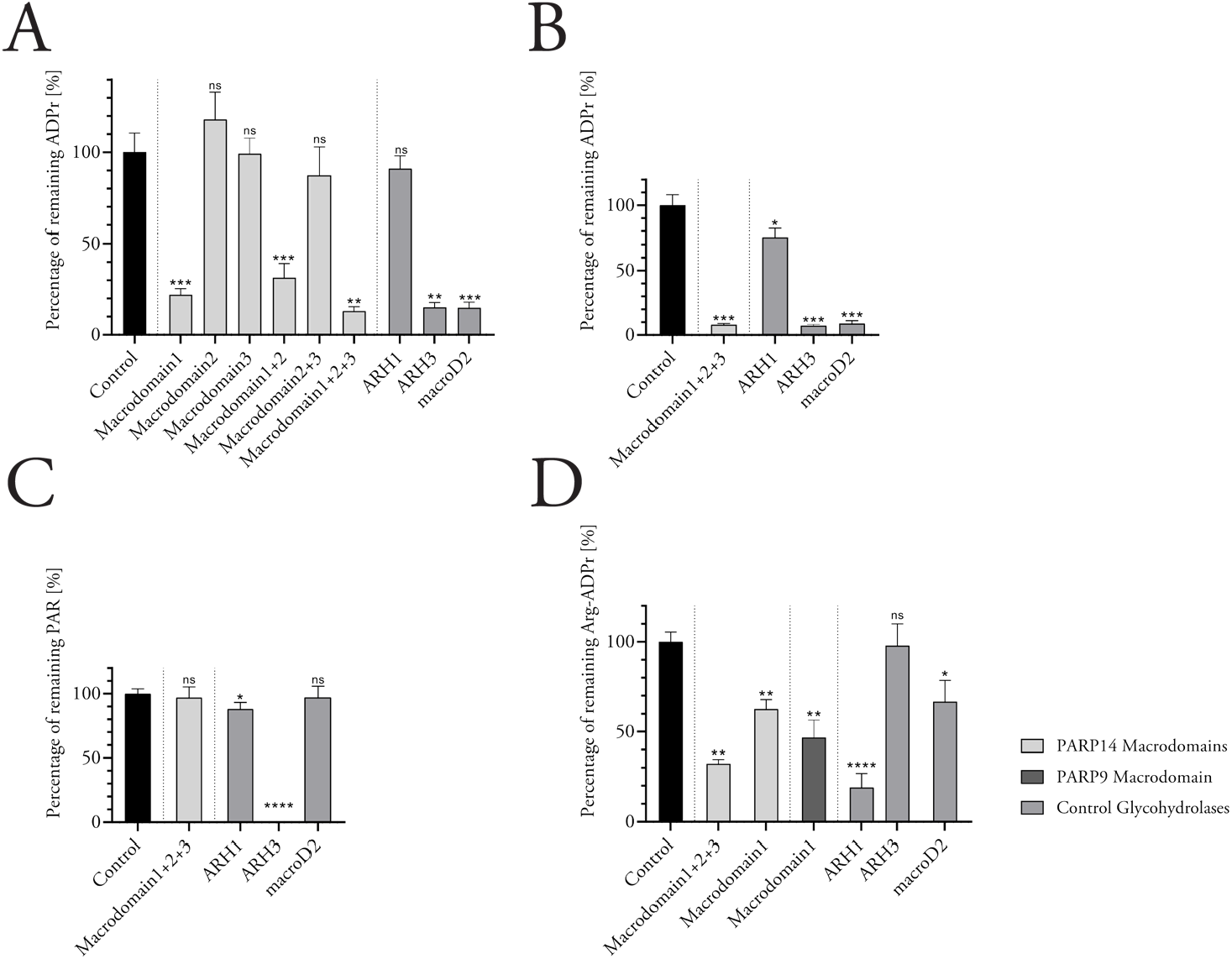
ADP-ribosyl glycohydrolase activity of PARP14 macrodomain-1 measured using a fluorescent protein overlay. (**A**) Percentage of ADP-ribosyl remaining on automodified PARP14 catalytic domain after treatment with macrodomains and known glycohydrolases as indicated. (**B**) Percentage of ADP-ribosyl remaining on automodified PARP10N819-V1007 after treatment with PARP14 macrodomain-1+2+3 and known glycohydrolases as indicated. (**C**) Percentage of PAR remaining on automodified PARP1 after treatment with PARP14 macrodomain-1+2+3 and glycohydrolases. (**D**) Percentage of ADP-ribosyl remaining on actin modified by C2I after treatment with PARP14 macrodomain-1 and macrodomain-1+2+3, PARP9 macrodomain 1 and glycohydrolases. Ann experiments; n=4; means ± S.D. are shown.

To further substantiate macrodomain-1 target preferences, we examined the processing of the catalytic domain of PARP10, which has previously been shown to be promiscuous in terms of target residues ((54) and references therein). The PARP14 macrodomain-1-2-3 construct as well as ARH3 and macroD2 all led to efficient removal of MARylation from PARP10 (Fig. 3B). Incubation of PARP10 with ARH1 resulted in a moderate, statistically significant reduction in MARylation levels. This confirmed our recent finding that Arg side chains are among the targets for PARP10 auto-MARylation (54). However, this result did not specifically answer the question of whether mac-rodomain-1 can process MARylated arginines. Therefore, we turned to actin, which is known to be ADP-ribosylated by the Clostridium botulinum C2I toxin subunit on a single arginine residue (Arg177) (55). Our analysis showed that PARP14 macrodomain-1 and PARP9 macrodomain-1 both elicited a strong and statistically significant reduction in actin ADP-ribosylation (Fig. 3D). ARH1 gave a near-complete removal of MAR signal, as expected, whereas ARH3 only slightly reduced MARylation levels on actin. These results strongly suggest that PARP14 and PARP9 macro-domains-1 are capable of cleaving ADP-ribosylated arginine.

Finally, we examined whether macrodomain-1 could cleave poly(ADP)ribose on PARP1, either on mixed sites or, when synthesized in the presence of HPF1, on Ser residues.(56) As shown in Fig. 3C and Fig. S5, only ARH3 could reduce PARylation under both conditions and did so efficiently, as expected (52).

### Catalytic mechanism of macrodomain-1 ADR-ribosyl glycohydrolase activity

Finally, we used site-directed mutagenesis to examine whether the active-site sequence similarities between macrodomain-1 and the Mac1 of the SARS-CoV and the SARS-CoV-2 Nsp3 protein also translated into similar catalytic mechanisms. We replicated several Mac1 mutants that resulted in reduced enzymatic activity in Mac1 (25,57,58). PARP14 and PARP9 macrodomain-1 carrying the mutations G924E and N940A (PARP14 sequence numbering) could not be purified owing to protein aggregation. Whether these mutations will permit soluble protein production when placed in a multidomain context remains to be examined. PARP14 macrodomain-1 mutant F926A and the homologous mutant PARP9 macrodomain-1 F244A were purified and both displayed substantially reduced catalytic activity (Fig. 4A,B). Although the mutations reduced the melting points of the domains, an excess of free ADP-ribose elevated the Tms of both mutant proteins to a similar degree as those of the wild type constructs (Fig. 4C). We conclude that these Phe side chains in PARP14 and PARP9 have similar roles as established for other ADP-ribosyl glycohydrolases, namely, to orient and/or stabilize the target ADP-ribosyl in the active site for catalysis. This is supported by the conservation of a small aromatic (Phe/ Tyr) side chain among catalytically active macrodomains but not among the PARP15 macrodomains in general (Fig. 5). Thus, while we present evidence for a flexible catalytic mechanism that can accommodate ADP-ribosyl cleavage off chemically different side chains, we have identified active-site mutations that strongly reduce macrodomain-1 in PARP14 and PARP9 and that will be useful to study the functions of these domains.

**Figure 4.**
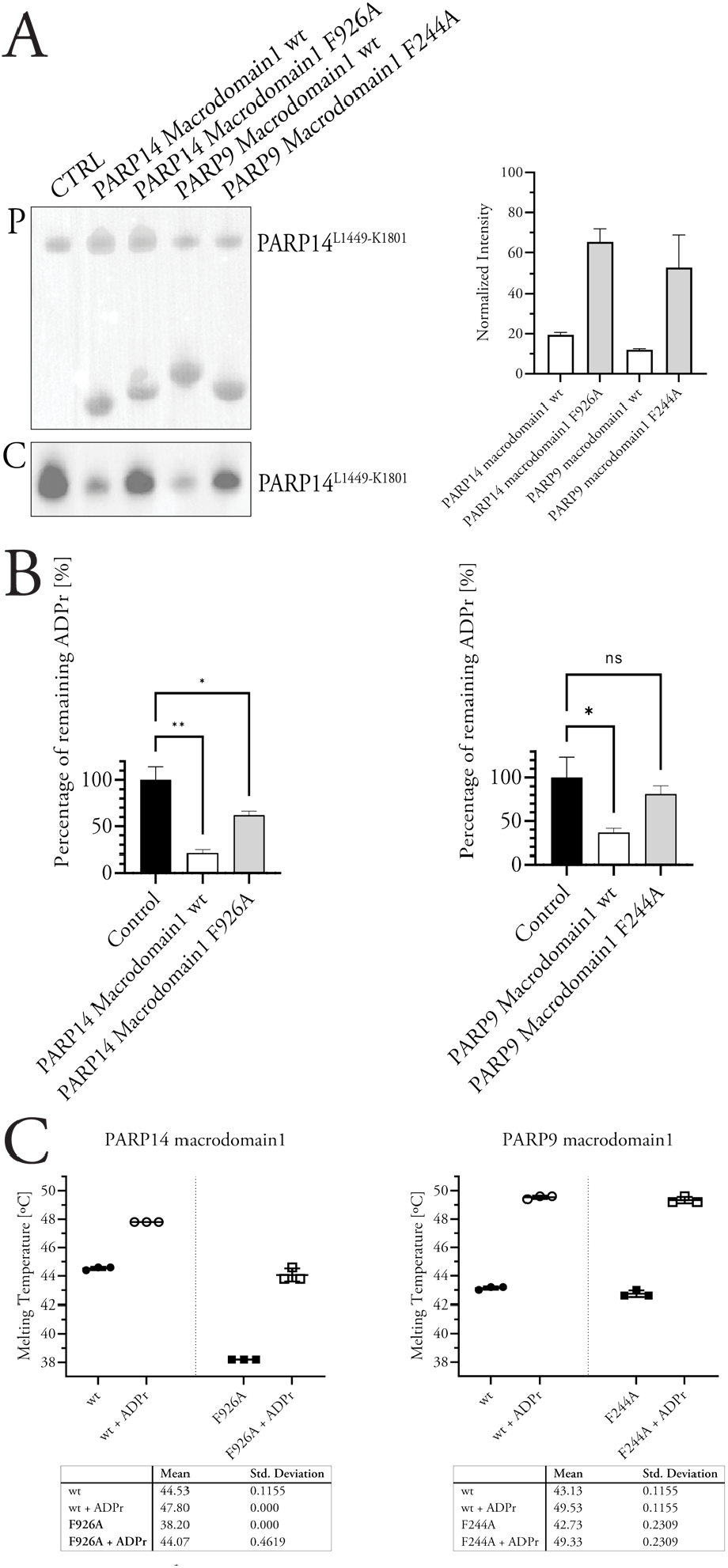
Mutagenesis of PARP14 and PARP9 macrodomain-1. (**A**) Automodified PARP14L1449-K1801 incubated with wild-type and mutant PARP14 and PARP9 macrodomain-1. “P”, Ponceau stained membrane; “C”, chemiluminescence. Normalized band intensities are shown on the right. (**B**) Reactions as in panel A were analyzed for remaining ADP-ribosylation levels using MacroGreen. (**C**) Melting temperature analysis using DSF for the wild type and mutant macrodomains in the absence and presence of free ADP-ribose.

**Figure 5.**
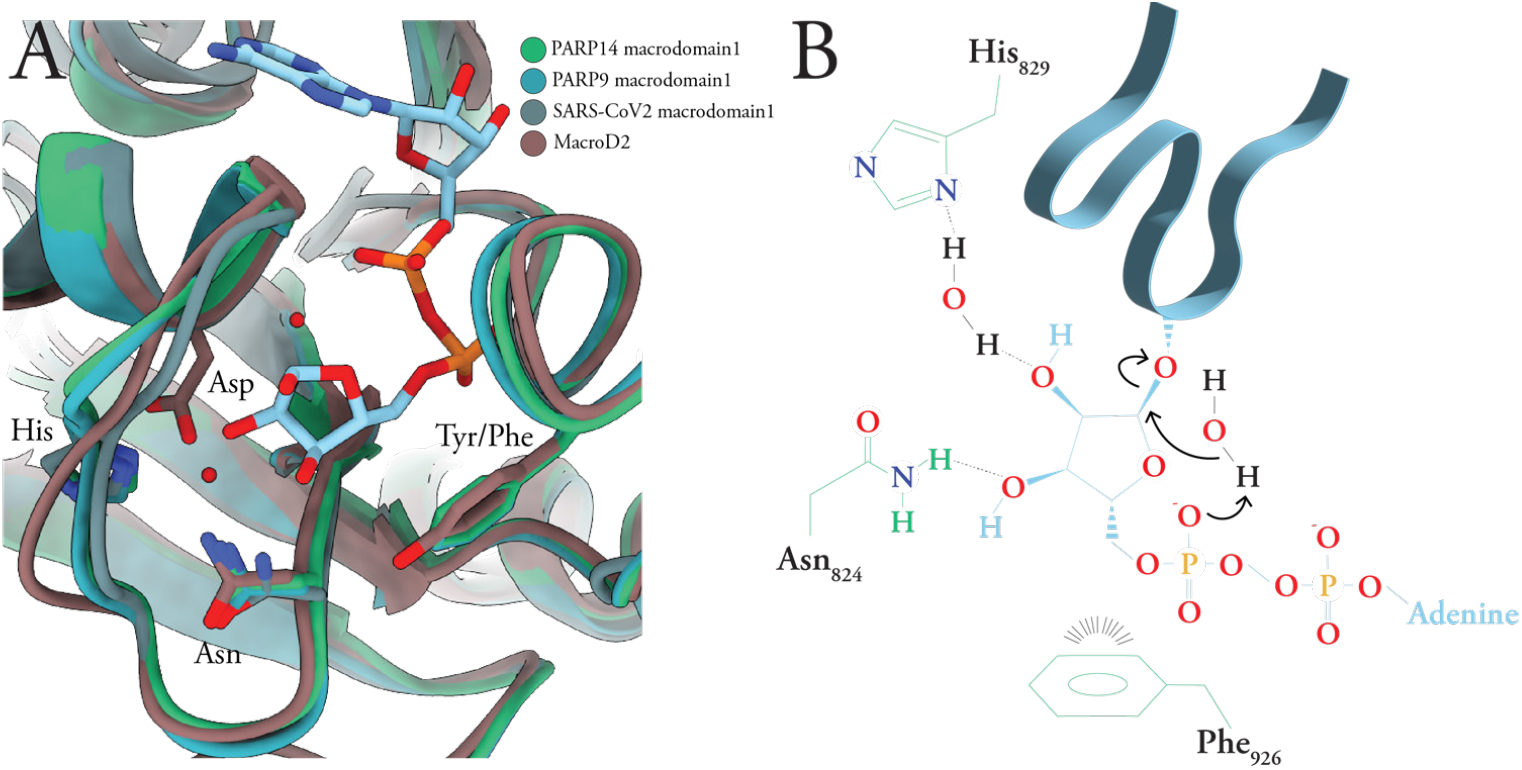
Details of ADP-ribose binding in the active site of macrodomain-1. (**A**) Superimposition PARP14 and PARP9 macrodomain-1, macroD2, and SARS-CoV-2 macrodomain-1 with the side chains discussed in the text and the ADP-ribose molecule bound to PARP14 shown as sticks. (**B**) Schematic illustration of a possible catalytic mechanism for PARP14 macrodomain-1, based on our results as well as (41,45-48).

### Conclusion and outlook

This study uncovered macrodomains-1 of PARP14 and the homologous PARP9 as novel ADP-ribosyl glycohydrolases. We show that PARP14 macrodomain-1 is a non-specific de-MARylating enzyme that can hydrolyze ADP-ribosyl off of Ser, Arg, and carboxylic acid side chains but cannot hydrolyze PAR chains. It remains to be seen whether macrodomain-1 can process MARylated nucleic acids as well. These findings will change the way we study PARP14 and PARP9 functions both in vitro and in cells. The Phe-to-Ala mutation that we showed to drastically reduce the activity of these enzymes might be a way to isolate macrodomain-1 functions from the functions of other domains. Pharmacological inhibitors of these macrodomains are clearly within reach and the compounds we identified here might serve as a starting point for the development of potent and cell-active marcodomain-1 inhibitors.

## Acknowledgments

We thank Drs. Emilia Strandback, Henry Ampah-Korsah and Tomas Nyman (Protein Science Facility at Karolinska Institutet, Stockholm) for molecular cloning. We also thank Drs. Johannes Rack and Ivan Ahel (Oxford University) for the ARH1 expression plasmid.

## Author Contributions

Conceptualization, A.T., C.C. and H.S.; investigation, A.T., C.C. and C.E.; formal analysis, A.T. and C.C.; writing— original draft preparation, A.T.; writing—review and editing, A.T. and H.S.; visualization, A.T.; supervision, H.S.; funding acquisition, H.S. All authors have read and agreed to the final version of the manuscript.

