## Supplemental Information Document for "PARP14 is a writer, reader and eraser of mono-ADP-ribosylation"

This file includes:

Supplementary Experimental Procedures
Supplementary Figure 1
Supplementary Figure 2
Supplementary Table 1
Supplementary Table 2
Supplementary Figure 3
Supplementary Figure 4
Supplementary Figure 5
Supplementary Figure 6

### Supplementary Experimental Procedures

#### Thermal stability and ligand induced Tm shift analysis by differential scanning fluorimetry

Melting point ($T_m$) assays were performed in white 96-well PCR plates (Bio-Rad #MLL9651). Every well contained a 25 µL solution of 0.4 mg/mL of enzyme, 2 mM of Z5010894420, Z5014193706, or Z5183357278 (Enamine), and SyproOrange (ThermoFisher; at 1:5000 dilution) in 50 mM HEPES pH 7.5, 300 mM NaCl, 10 %v/v glycerol, 0.5 mM TCEP buffer. Control wells contained either 0, 2, or 8% DMSO. Plates were incubated at 20 °C for 10 seconds and then the temperature was increased by 1 °C/min up to 95 °C. Fluorescence signals were measured with the CFX96 Touch Real-Time PCR Detection System (Bio-Rad) and data analysis was performed with the CFX Manager software (version 3.1, Bio-Rad).

PARP14 macrodomain-1 compound inhibition assayADP-ribosyltaion reactions were performed as described in the main text. 50 µL per well of solutions of PARP14 macrodomain-1 at a final concentration of 3 µM and different concentrations (10, 50, 100, 500 and 1000 µM) of SARS-CoV-2 Nsp3 Mac1 inhibitors (Enamine) or 1, 2, or 4% DMSO were added, except for positive control (omission of macrodomain), negative control (omission of NAD+) and blank, where 50 µL of 50 mM HEPES pH 7.5, 100 mM NaCl, 0.2 mM TCEP, 4 mM MgCl2, 0.1 mM EDTA buffer was added. The plates processed and analyzed as stated in the main text.

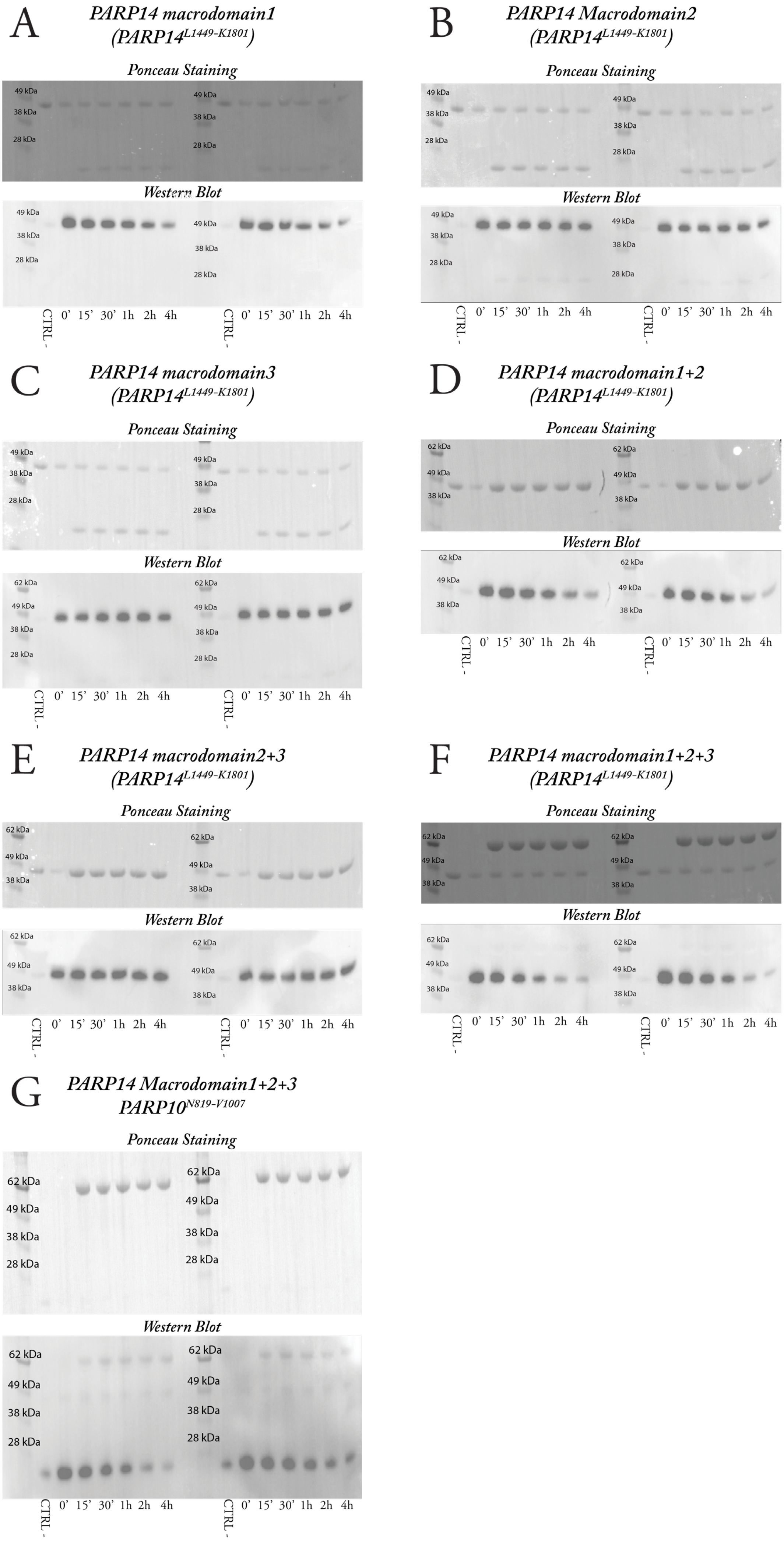
A
PARP14 macrodomain1
(PARP14L1449-K1801)
Ponceau Staining
Western Blot
B
PARP14 Macrodomain2
(PARP14L1449-K1801)
Ponceau Staining
Western Blot
C
PARP14 macrodomain3
(PARP14L1449-K1801)
Ponceau Staining
Western Blot
D
PARP14 macrodomain1+2
(PARP14L1449-K1801)
Ponceau Staining
Western Blot
E
PARP14 macrodomain2+3
(PARP14L1449-K1801)
Ponceau Staining
Western Blot
F
PARP14 macrodomain1+2+3
(PARP14L1449-K1801)
Ponceau Staining
Western Blot
G
PARP14 Macrodomain1+2+3
PARP10N819-V1007
Ponceau Staining
Western Blot
62 kDa
49 kDa
38 kDa
28 kDa
CTRL -
0' 15' 30' 1h 2h 4h

#### Previous page Figure S1 (related to Fig. 1 of the main text)

**A-G** Original membranes, including duplicate reactions and molecular weight markers, of the experiments shown in Fig. 1 of the main text. Experiments not shown in the main text are those pertaining PARP14 macrodomain-2 and -3 (panels B and C, respectively).

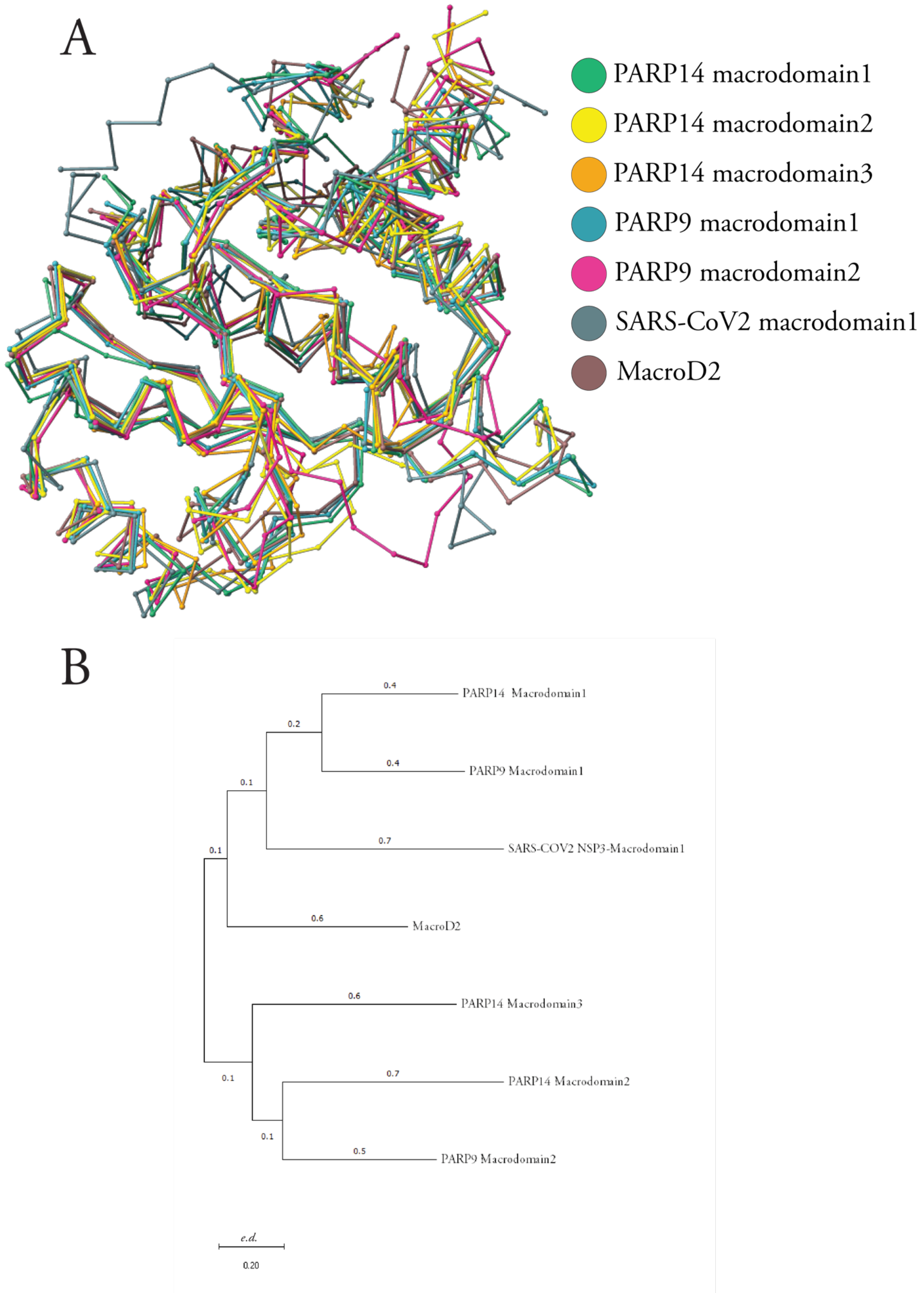


#### Figure S2 (related to Fig. 2 of the main text)

**A** Alignment of the crystal structures of the macrodomains shown in the sequence alignment of Fig.1B of the main text. **B** Phylogenetic tree of the same sequences calculated using MEGA11.

***Table 1. Identity and similarity scores.***

| | PARP14 macrodomain1 | PARP9 macrodomain1 | PARP14 macrodomain2 | PARP9 macrodomain2 | PARP14 macrodomain3 | macroD2 | NSP3 macrodomain1 | |
|---|---|---|---|---|---|---|---|---|
| **PARP14 macrodomain1** | - | 54.25% | 33.51% | 29.67% | 35.46% | 37.79% | 40.65% | Similarity |
| **PARP9 macrodomain1** | 42.02% | - | 34.04% | 28.57% | 33.72% | 37.20% | 40.65% | |
| **PARP14 macrodomain2** | 22.87% | 21.8% | - | 43.95% | 34.88% | 23.25% | 32.41% | |
| **PARP9 macrodomain2** | 15.93% | 17.03% | 31.31% | - | 40.69% | 25.58% | 32.41% | |
| **PARP14 macrodomain3** | 21.51% | 19.18% | 22.67% | 31.39% | - | 25.00% | 33.72% | |
| **macroD2** | 23.83% | 26.16% | 15.69% | 18.02% | 14.53% | - | 34.30% | |
| **NSP3 macrodomain1** | 29.12% | 26.37% | 20.32% | 23.62% | 23.83% | 22.09% | - | |
| | | | | Identity | | | | |

***Table 2. Cα RMSD values* calculated from the structural alignment in Figure 1 B.***

| | PARP14 macrodomain2 | PARP14 macrodomain3 | PARP9 macrodomain1 | PARP9 macrodomain2 | NSP3 macrodomain1 | macroD2 |
|---|---|---|---|---|---|---|
| **PARP14 macrodomain1** | 1.460 | 1.333 | 0.546 | 1.705 | 1.331 | 0.721 |
| **§PARP14 macrodomain2** | - | 1.243 | 1.220 | 1.055 | 4.328 | 1.167 |
| **PARP14 macrodomain3** | - | - | 1.187 | 1.097 | 6.415 | 1.194 |
| **PARP9 macrodomain1** | - | - | - | 1.146 | 1.968 | 0.798 |
| **PARP9 macrodomain2** | - | - | - | - | 1.346 | 1.321 |
| **NSP3 macrodomain1** | - | - | - | - | - | 1.832 |

*Alignment RMSD values are expressed in Ångstrom

A

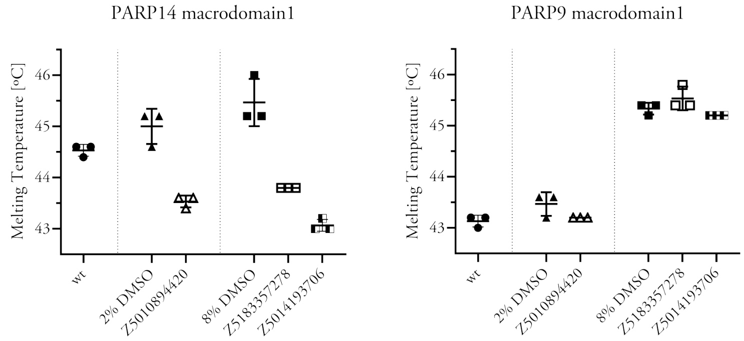


| | Mean | Std. Deviation |
|---|---|---|
| wt | 44.53 | 0.1155 |
| 2% DMSO | 45.00 | 0.3464 |
| Z5010894420 | 43.53 | 0.1155 |
| 8% DMSO | 45.47 | 0.4619 |
| Z5183357278 | 43.80 | 0.000 |
| Z5014193706 | 43.07 | 0.1155 |

| | Mean | Std. Deviation |
|---|---|---|
| wt | 43.13 | 0.1155 |
| 2% DMSO | 43.47 | 0.2309 |
| Z5010894420 | 43.20 | 0.000 |
| 8% DMSO | 45.33 | 0.1155 |
| Z5183357278 | 45.53 | 0.2309 |
| Z5014193706 | 45.20 | 0.000 |

B

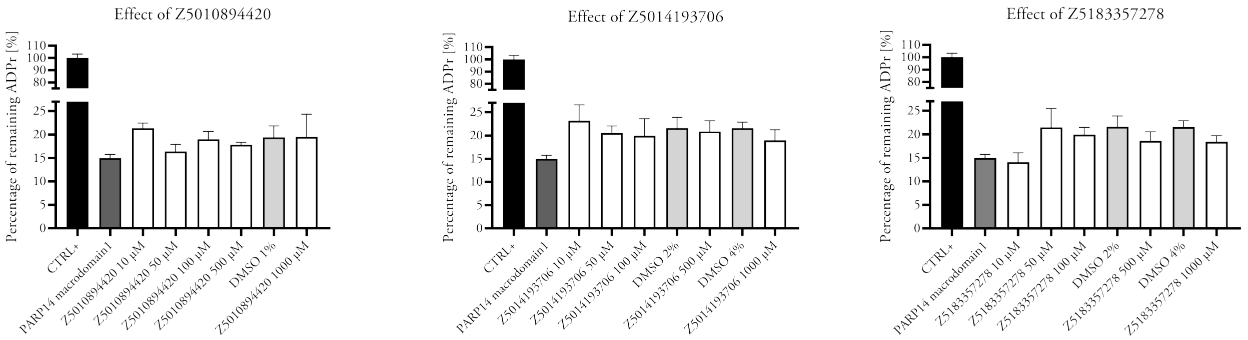


#### Figure S3

Effect of SARS-CoV-2 Nsp3 Mac1 inhibitors on PARP14 and PARP9 macrodomain-1. **A** Melting point shift assays of PARP14 (left) respectively PARP9 (right) macrodomain-1 in the presence of 2 mM of the indicated compounds. The effect of DMSO, at the concentrations required for this concentration for the respective compounds, is also shown. All three compounds destabilized PARP14 macrodomain-1 and had no effect on the Tm of PARP9 macrodomain-1. **B** Inhibition assays (using detection of remaining ADP-ribosylation levels using MacroGreen) of PARP14 macrodomain-1 activity over automodified PARP14. The relevant concentrations of DMSO are shown in light grey bars. None of the compounds significantly inhibited macrodomain-1.

#### Next page Figure S4 (related to Fig. S3)

Effect of SARS CoV 2 Nsp3 Mac1 inhibitors on PARP14 and PARP9 macrodomain-1. . Left panels, thermal melting profiles; right hand panels, first derivatives.

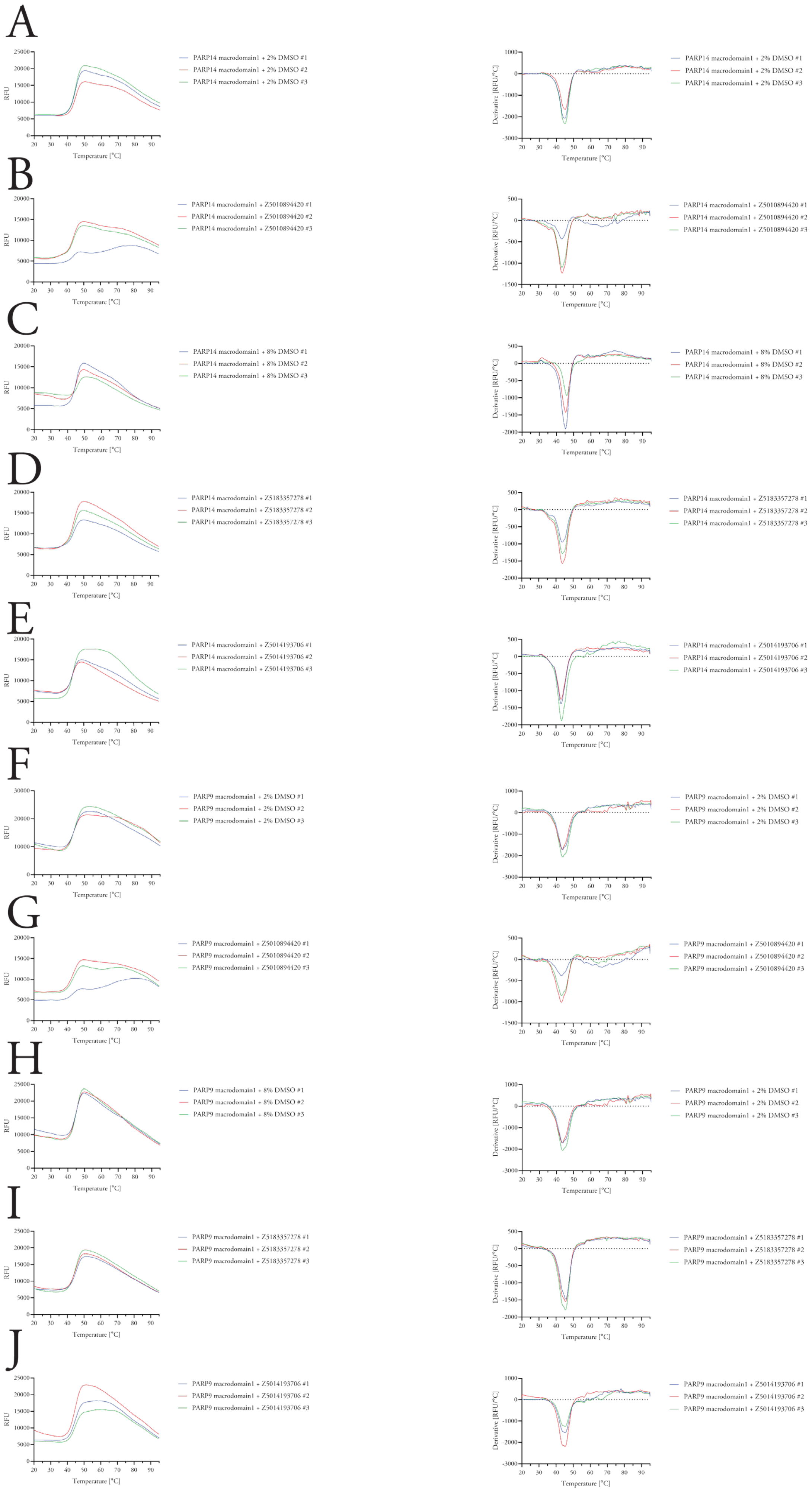
A
RFU
Temperature [°C]
PARP14 macrodomain1 + 2% DMSO #1
PARP14 macrodomain1 + 2% DMSO #2
PARP14 macrodomain1 + 2% DMSO #3
Derivative [RFU/°C]
Temperature [°C]
PARP14 macrodomain1 + 2% DMSO #1
PARP14 macrodomain1 + 2% DMSO #2
PARP14 macrodomain1 + 2% DMSO #3
B
PARP14 macrodomain1 + Z5010894420 #1
PARP14 macrodomain1 + Z5010894420 #2
PARP14 macrodomain1 + Z5010894420 #3
PARP14 macrodomain1 + Z5010894420 #1
PARP14 macrodomain1 + Z5010894420 #2
PARP14 macrodomain1 + Z5010894420 #3
C
PARP14 macrodomain1 + 8% DMSO #1
PARP14 macrodomain1 + 8% DMSO #2
PARP14 macrodomain1 + 8% DMSO #3
PARP14 macrodomain1 + 8% DMSO #1
PARP14 macrodomain1 + 8% DMSO #2
PARP14 macrodomain1 + 8% DMSO #3
D
PARP14 macrodomain1 + Z5183357278 #1
PARP14 macrodomain1 + Z5183357278 #2
PARP14 macrodomain1 + Z5183357278 #3
PARP14 macrodomain1 + Z5183357278 #1
PARP14 macrodomain1 + Z5183357278 #2
PARP14 macrodomain1 + Z5183357278 #3
E
PARP14 macrodomain1 + Z5014193706 #1
PARP14 macrodomain1 + Z5014193706 #2
PARP14 macrodomain1 + Z5014193706 #3
PARP14 macrodomain1 + Z5014193706 #1
PARP14 macrodomain1 + Z5014193706 #2
PARP14 macrodomain1 + Z5014193706 #3
F
PARP9 macrodomain1 + 2% DMSO #1
PARP9 macrodomain1 + 2% DMSO #2
PARP9 macrodomain1 + 2% DMSO #3
PARP9 macrodomain1 + 2% DMSO #1
PARP9 macrodomain1 + 2% DMSO #2
PARP9 macrodomain1 + 2% DMSO #3
G
PARP9 macrodomain1 + Z5010894420 #1
PARP9 macrodomain1 + Z5010894420 #2
PARP9 macrodomain1 + Z5010894420 #3
PARP9 macrodomain1 + Z5010894420 #1
PARP9 macrodomain1 + Z5010894420 #2
PARP9 macrodomain1 + Z5010894420 #3
H
PARP9 macrodomain1 + 8% DMSO #1
PARP9 macrodomain1 + 8% DMSO #2
PARP9 macrodomain1 + 8% DMSO #3
PARP9 macrodomain1 + 2% DMSO #1
PARP9 macrodomain1 + 2% DMSO #2
PARP9 macrodomain1 + 2% DMSO #3
I
PARP9 macrodomain1 + Z5183357278 #1
PARP9 macrodomain1 + Z5183357278 #2
PARP9 macrodomain1 + Z5183357278 #3
PARP9 macrodomain1 + Z5183357278 #1
PARP9 macrodomain1 + Z5183357278 #2
PARP9 macrodomain1 + Z5183357278 #3
J
PARP9 macrodomain1 + Z5014193706 #1
PARP9 macrodomain1 + Z5014193706 #2
PARP9 macrodomain1 + Z5014193706 #3
PARP9 macrodomain1 + Z5014193706 #1
PARP9 macrodomain1 + Z5014193706 #2
PARP9 macrodomain1 + Z5014193706 #3

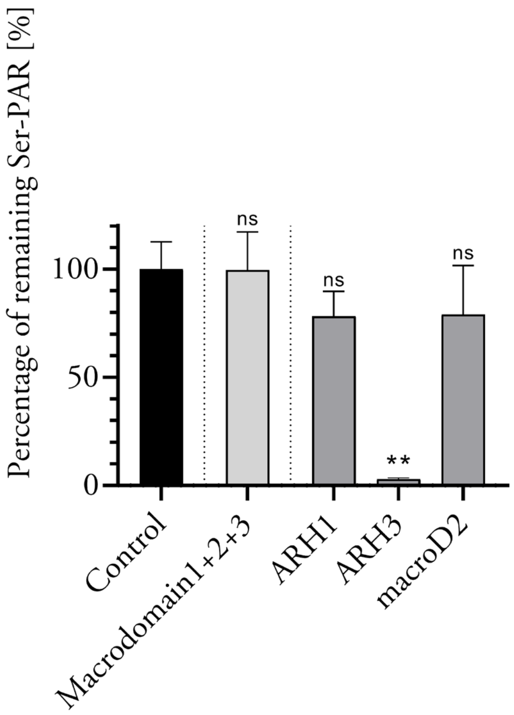


#### Figure S5 (related to Fig. 3 of the main text)

Quantification using MacroGreen of ADP-ribosylation levels present after incubation of PARP1 with HPF1 and NAD+ and subsequent incubation with the indicated macrodomains. Error bars represent S.D. (n=4) and significance levels refer to the control.

#### Next page Figure S6 (related to Fig. 4 of the main text)

Differential Scanning Fluorimetry (DSF) of PARP14 and PARP9 macrodomain-1 wild type and F->A mutants in presence and absence of free ADP-ribose. Left panels, thermal melting profiles; right hand panels, first derivatives.

A

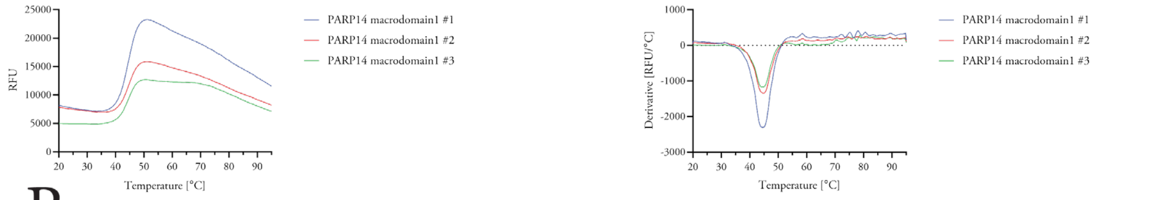
RFU
Temperature [°C]
PARP14 macrodomain1 #1
PARP14 macrodomain1 #2
PARP14 macrodomain1 #3
Derivative [RFU/°C]
Temperature [°C]
PARP14 macrodomain1 #1
PARP14 macrodomain1 #2
PARP14 macrodomain1 #3

B

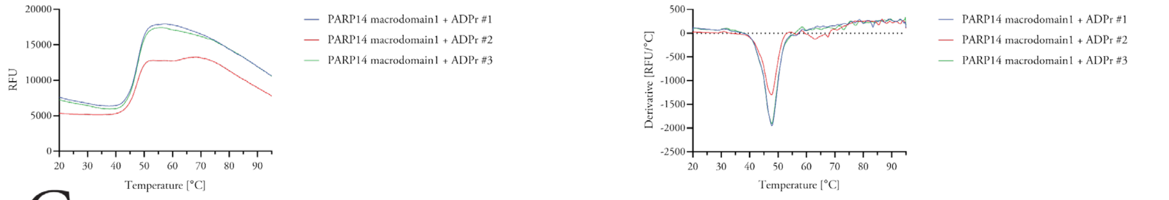
RFU
Temperature [°C]
PARP14 macrodomain1 + ADPr #1
PARP14 macrodomain1 + ADPr #2
PARP14 macrodomain1 + ADPr #3
Derivative [RFU/°C]
Temperature [°C]
PARP14 macrodomain1 + ADPr #1
PARP14 macrodomain1 + ADPr #2
PARP14 macrodomain1 + ADPr #3

C

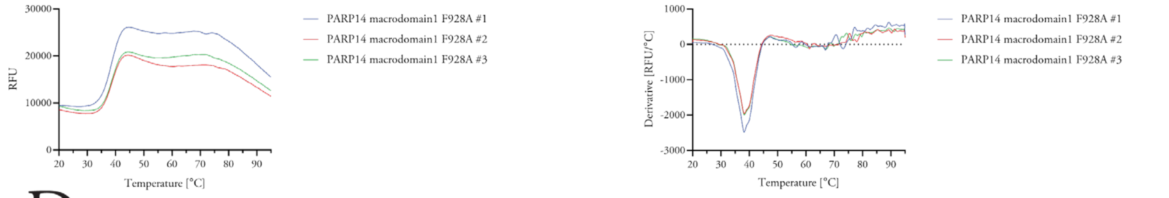
RFU
Temperature [°C]
PARP14 macrodomain1 F928A #1
PARP14 macrodomain1 F928A #2
PARP14 macrodomain1 F928A #3
Derivative [RFU/°C]
Temperature [°C]
PARP14 macrodomain1 F928A #1
PARP14 macrodomain1 F928A #2
PARP14 macrodomain1 F928A #3

D

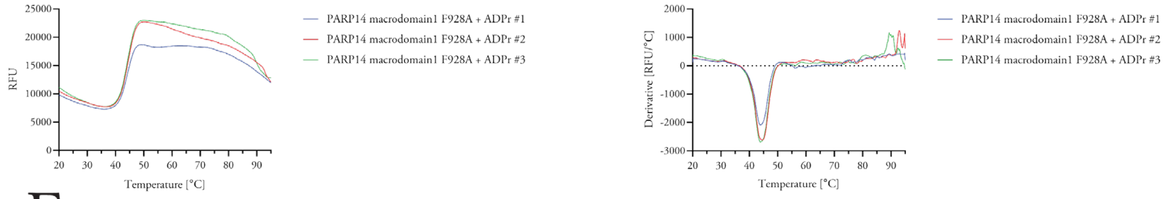
RFU
Temperature [°C]
PARP14 macrodomain1 F928A + ADPr #1
PARP14 macrodomain1 F928A + ADPr #2
PARP14 macrodomain1 F928A + ADPr #3
Derivative [RFU/°C]
Temperature [°C]
PARP14 macrodomain1 F928A + ADPr #1
PARP14 macrodomain1 F928A + ADPr #2
PARP14 macrodomain1 F928A + ADPr #3

E

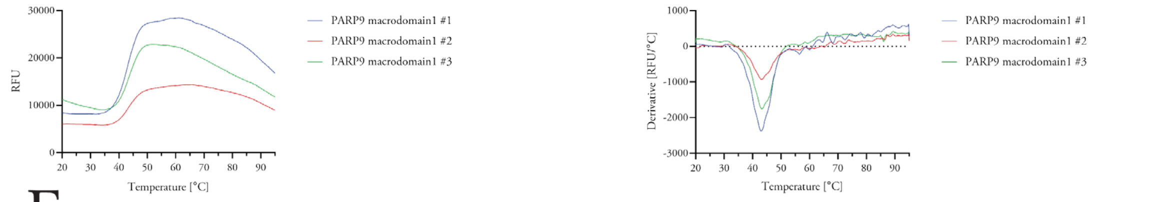
RFU
Temperature [°C]
PARP9 macrodomain1 #1
PARP9 macrodomain1 #2
PARP9 macrodomain1 #3
Derivative [RFU/°C]
Temperature [°C]
PARP9 macrodomain1 #1
PARP9 macrodomain1 #2
PARP9 macrodomain1 #3

F

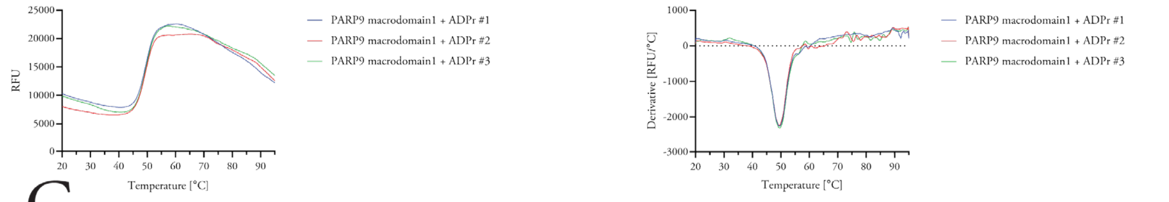
RFU
Temperature [°C]
PARP9 macrodomain1 + ADPr #1
PARP9 macrodomain1 + ADPr #2
PARP9 macrodomain1 + ADPr #3
Derivative [RFU/°C]
Temperature [°C]
PARP9 macrodomain1 + ADPr #1
PARP9 macrodomain1 + ADPr #2
PARP9 macrodomain1 + ADPr #3

G

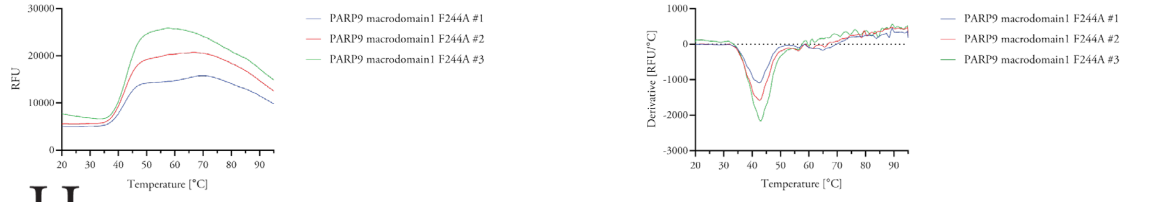
RFU
Temperature [°C]
PARP9 macrodomain1 F244A #1
PARP9 macrodomain1 F244A #2
PARP9 macrodomain1 F244A #3
Derivative [RFU/°C]
Temperature [°C]
PARP9 macrodomain1 F244A #1
PARP9 macrodomain1 F244A #2
PARP9 macrodomain1 F244A #3

H

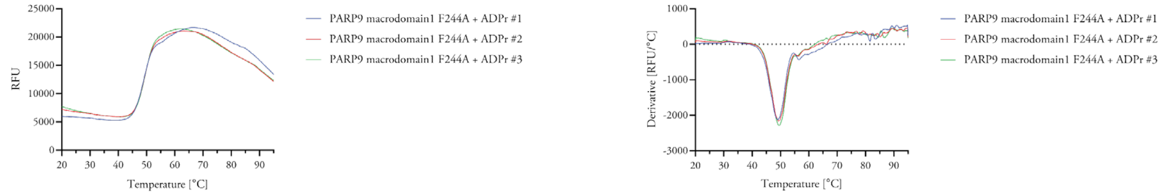
RFU
Temperature [°C]
PARP9 macrodomain1 F244A + ADPr #1
PARP9 macrodomain1 F244A + ADPr #2
PARP9 macrodomain1 F244A + ADPr #3
Derivative [RFU/°C]
Temperature [°C]
PARP9 macrodomain1 F244A + ADPr #1
PARP9 macrodomain1 F244A + ADPr #2
PARP9 macrodomain1 F244A + ADPr #3
